# Structure-function relationship of PE11 esterase of *Mycobacterium tuberculosis* with respect to its role in virulence

**DOI:** 10.1101/2024.06.18.599654

**Authors:** Priyanka Dahiya, Amit Banerjee, Abhishek Saha, Vinay Kumar Nandicoori, Sudip Ghosh, Sangita Mukhopadhyay

## Abstract

The lipolytic enzymes of *Mycobacterium tuberculosis* play a critical role in immunomodulation and virulence. Among these proteins, PE11 which also belongs to the PE/PPE family, is the smallest (∼10.8 kDa) and play a significant role in cell wall remodelling and virulence. PE11 is established to be an esterase, but its enzymatic and structural properties are not yet characterized. In this study, using homology modelling we deduced the putative structure which shows the presence of both α-helix and β-sheet structures which is in close agreement with that observed by CD spectra of the purified protein. PE11 was found to contain a GX3SX4G motif homologous to canonical ‘GxSxG’ motif present in many serin hydrolases. The catalytic triad appears to be located within this motif as substitution of Serine^26^ and Glycine^31^ residues abrogated its enzymatic activity. Gel-filtration chromatography data indicate that PE11 possibly exists as dimer and tetramer showing positive cooperativity for binding its substrates. In addition, PE11 esterase activity was found to be critical for cell wall remodelling, antibiotic resistance and conferring survival advantages to *M. tuberculosis*. Our data suggest that PE11 can be targeted for designing potential therapeutic strategies.

## Introduction

Tuberculosis remained a constant global health challenge, causing an estimated 1.4 million deaths annually (World Health Organization TB Report, 2023). Whole genome sequencing of *Mycobacterium tuberculosis* (M.tb) unravelled known or putative functions of most of the genes, but function of about 16% genes remained unclear (Cole et al., 1998). This includes the PE (Pro-Glu)/PPE (Pro-Pro-Glu) gene family, which constitutes approximately 10% of the M.tb genome of which 99 PE genes were found to be present in a laboratory strain *Mycobacterium tuberculosis* (*H37Rv)* (Sampson et al., 2011; Brennan and Delogu, 2002). The PE family proteins are characterized by the presence of a conserved N-terminal domains and have been implicated to be involved with M.tb immunogenicity (Delogu and Brennan, 2001; Deng et al., 2017). In addition, various studies suggest their involvement in delivering C-terminal cargo to the type VII secretion systems (Santucci et al., 2019).

Insights into the bacterium’s prolonged survival within the host has highlighted the crucial roles of several lipases and esterases in lipid metabolism, nutrient accumulation, cell wall remodelling, intracellular survival, pathogenesis, and the host’s immune invasion (Singh et al., 2016; Yang et al., 2019). To date, among all the PE proteins, only PE1, PE2, PE11 and PE16, have been reported to possess lipase/esterase activity (Sultana et al., 2013, Divya et al., 2018; Singh et al., 2016). Among these, PE1 and PE2 were found to contain a conserved ‘α/β hydrolase’ domain, characteristic of α/-β serine hydrolase family of esterases and are co-transcribed from the same operon with substrate specificity for short to medium chain (C2- C8) acyl esters (Divya et al., 2018). On the other hand, PE16 exhibited a notable specificity for the 6-carbon chain fatty acid ester substrate (Sultana et al., 2013). In addition, PE11 protein was found to play a role in cell wall remodelling with a preference for shorter chain (C2-C4) acyl esters (Singh et al., 2016).

The intricate interplay between lipid metabolism and the virulence of various intracellular bacteria is widely recognized. The lipolytic enzymes play a crucial function by breaking down host cell lipids and releasing free fatty acids which not only serve as essential energy sources but also act as fundamental building blocks for synthesizing cell envelope components. Moreover, the released fatty acids also contribute to the modulation of host immune responses, thereby influencing the pathogenicity of the bacteria within the host environment (Rameshwaram et al., 2018). In this context, the Lip Family of M.tb, comprising 24 members characterized by α/β-hydrolase, plays a vital role. These enzymes exhibit remarkable diversity in enzymatic function, active site sequence, and structure (Bauer et al., 2020). Numerous studies have explored the lipase/esterase activity of various proteins of the Lip family, particularly Rv3775 (LipE), LipL, and LipD, and highlighted their roles in virulence and the host’s innate and adaptive immune responses. Also, the enhanced pathogenicity observed with the overexpression of LipY in *M. tuberculosis* H37Rv is linked to the suppression of the host’s protective immune response. In addition, presence of LipL protein was found to induce a strong humoral immune response in humans and triggered a CD8^+^ T cell-mediated response in mice (Yang et al., 2019; Cao et al., 2015; Singh et al., 2014).

PE11 (Rv1169c), also known as LipX is part of both PE as well as Lip family of proteins. PE11 is upregulated significantly under varied stress conditions. The protein is known for its association with the modification of cell wall architecture and conferring survival advantage to *Mycobacteria* (Betts et al., 2002; Fisher et al., 2002; Rustad et al., 2008; Singh et al., 2016; Rastogi et al. 2017). It has been reported that the PE11 protein plays a crucial role during *ex vivo* and *in vivo* infection (Singh et al., 2016; Rastogi et al., 2017). Herein, we aim to further understand its functional significance in mycobacterial pathophysiology by enzymatic characterization and identification of the active site pocket, including critical amino acids. Moreover, our study emphasizes the crucial role of PE11’s esterase activity, not only in facilitating bacterial growth *in vitro* but also in its involvement during infection within macrophages. These findings further highlight the functional significance of the PE11 protein in mycobacterial virulence and suggest that PE11 as a promising therapeutic target for tuberculosis.

## Materials and methods

### Purification of wild-type recombinant PE11

The wild-type PE11 sequence (100 amino acids) with His-tag at the C-terminal was cloned in the pET23a vector and the existence of the insert was confirmed by restriction digestion followed by sequencing. The clones were then transformed into *Escherichia coli* BL21(DE3) bacterium. Transformed bacteria were further cultured in Terrific Broth containing 100 mg/ml ampicillin at 37°C. For induction, Isopropyl 1-thio-β-D-galactopyranoside (VWR Amresco, USA) was added to the mid-log phase culture at a concentration of 0.1 mM. His-tagged PE11 recombinant protein was purified using Ni-NTA metal affinity resin (Clontech Laboratories, USA) in denaturing conditions using 8 M urea (Sigma-Aldrich, USA); the protein was refolded back to its native state by sequentially removing the urea in a stepwise manner. The recombinant PE11 was stored in a Tris-HCl-based buffer.

### Bacterial culture

*E. coli* bacteria were grown in Luria-Bertani broth (BD Difco^TM^). Mycobacterial strains were cultured in Middlebrook 7H9 medium (BD Difco^TM^) supplemented with 10% albumin, dextrose, NaCl, catalase (ADC) along with 0.2% glycerol (Sigma), and 0.05% Tween-80 (Sigma), or 7H10/7H11 agar (BD Difco^TM^) with 10% OADC (HiMedia) and 0.2% glycerol.

### Generation of various *M. smegmatis* strains

To facilitate site-directed mutagenesis of Rv1169c, we devised internal primers and conducted overlap PCR employing three distinct primer sets (see Table 1). These primers were specifically crafted to substitute S26, and G31 with Alanine within the wild-type PE11 sequence in pVV-16 plasmid, with the ultimate aim of enabling its transformation into *Mycobacterium smegmatis* (Msmeg).

**Table 1:**
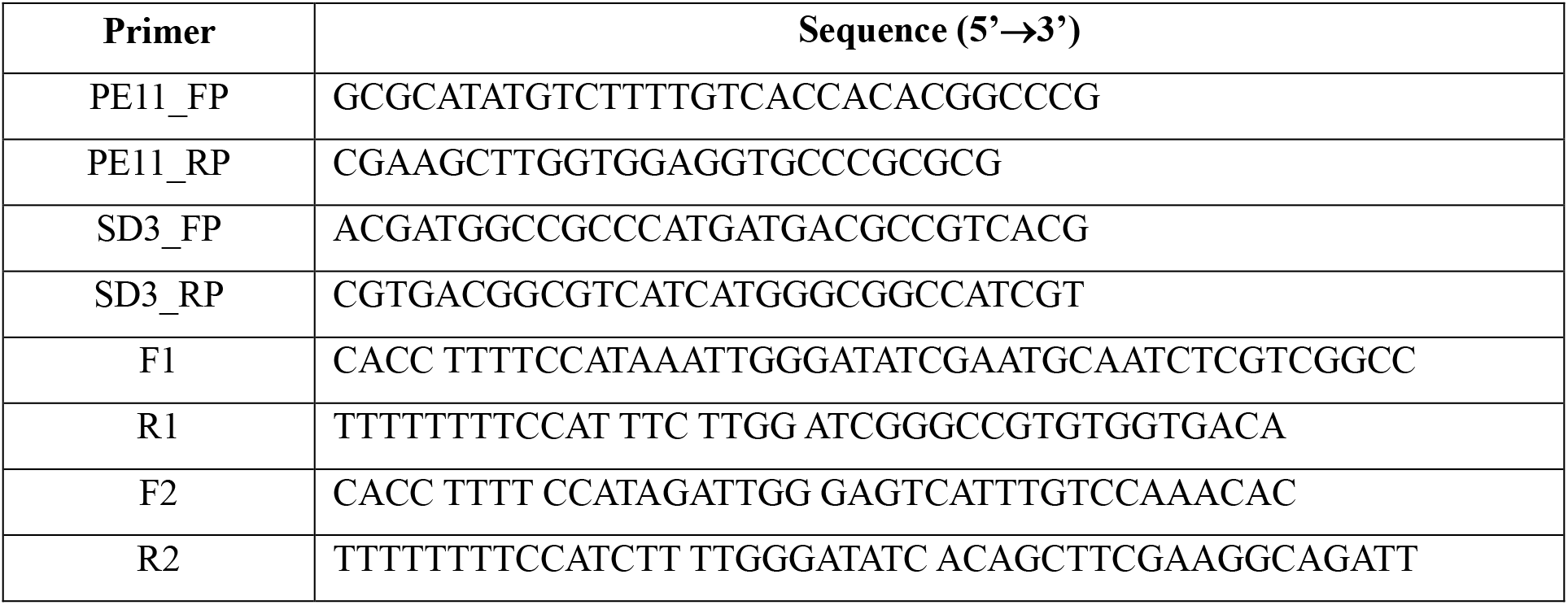
Primer list used for overlap PCR experiment.

### Generation of knock-out and complementary strains of *M. tuberculosis*

PE11 H37Rv knock-out (H37Rv-PE11KO) strain was generated using phage-mediated recombineering (van Kessel and Hatfull, 2007). Approximately 850 bp regions upstream (LHS) and 950 bp downstream (RHS) of the gene of interest including 30 bp within the gene of interest. The fragments were amplified *H37Rv* genomic DNA using specific primers (See T**able 1**). The fragments were digested and ligated with compatible ends oriE and the Hygromycin resistance gene (Hyg^r^) to construct an allelic exchange substrate. The AES was digested with Pac1 to excise the required fragment having LHS-Hyg^r^-RHS to be inserted in phAE159 phagemid. The Phage was generated using MaxPlax^TM^ and electroporated later. For generating complementary and mutant strains, a theophylline-based inducible riboswitch vector containing wild-type PE11 or mutant PE11(S3) sequence (S26 and G31 residues were replaced with Alanine), was electroporated into competent cells of *M. smegmatis* or H37Rv- PE11KO M.tb strain.

### Enzymatic Assay

For estimating the enzyme activity of PE11 protein, a broad pH ranges from pH 3.0 to pH 10.0 and a temperature range of 18°C to 47°C was considered. Enzyme activity was determined spectrophotometrically by hydrolysis of chromogenic substrates like p-Nitrophenol acetate (pNP-acetate) and p-Nitrophenol butyrate (pNP-butyrate) (Sigma-Aldrich) as described by Singh et al., (2016). For each assay (200 µl), 10 mM pNP-acetate or pNP-butyrate (Sigma-Aldrich, USA) dissolved in acetonitrile (ACN) and 100 mM Tris-HCl (pH 8.0) containing recombinantly purified PE11 protein (10 μg) were used. Incubation was carried out for 5 min at 30°C, and kinetics was measured at 405 nm for 20 min. The enzymatic hydrolysis was quantified by recording the absorbance of the hydrolyzed products of pNP-acetate/pNP-butyrate using SpectraMax M5 Multi-Mode Microplate Reader using a standard pNP calibration curve. The activity was expressed in international units (U, corresponding to 1 μmol of pNP released per minute).

### Circular Dichroism spectroscopy analysis

We used the purified His-tagged PE11 protein at a concentration of 0.30 mg/mL for Circular Dichroism (CD) spectroscopy. This protein was solubilized in a 20 mM Phosphate buffer with a pH of 8 and measurements were conducted at room temperature. We also investigated the changes in the enzyme’s secondary structure across a pH range (pH 4 to pH 12) within the Phosphate buffer system. The UV CD spectra of the protein were recorded between 190 and 250 nm using a Jasco J-810 Spectropolarimeter^™^, which was maintained at 25°C. The protein samples were contained in 1 mm quartz cuvettes, each containing 300 µl of the protein dissolved in the appropriate working buffer solutions. The spectroscopy data was collected with a data pitch of 0.1 nm, a bandwidth of 2.0 nm, and a scanning speed of 50 nm/min. The collected data were subsequently analyzed using the Jasco software^©^.

### Protein structure prediction

A restraint-based homology modelling approach was adopted for developing all the theoretical structures using MODELLER^©^ 10.0 (Webb and Sali, 2017; Webb and Sali, 2021). The analyses were done for PE11 and PE11(S3) with substituted mutations (S26A and G31A) sequences, each having 100 amino acid length. The template was searched using PSI-BLAST (Altschul et al., 1997; Jones and Swindells, 2002) based on the sequence identity as well as structural similarity with the wild-type PE11 sequence. In addition to the BLAST-based template search, the evaluation of plausible suitable templates was done using the Modbase (Pieper et al., 2002) and other comparative protein modelling approaches. Based on the presence of beta sheets (∼15%) identified through our CD analysis in PE11, we considered using the templates (PDB IDs: 1cex.pdb, 2czq.pdb, 3aja.pdb and 3hc7.pdb) as described by Sultana et al., (2011).

### Structure prediction and evaluation

All the 4 templates mentioned above were analyzed using a weighted pair group average clustering-based distance matrix, Modeller Objective Function (MOF) (Sánchez and Šali, 1997), Discrete Optimized Protein Energy (DOPE) (Shen and Sali, 2006), and GA341 scores (Eswar et al., 2008), and Ramachandran plot (Hooft et al., 1997), and were used for the multi template-based modelling. For each target protein, 5 models were developed and compared to obtain the best model. Structure with lowest MOF was considered for further analyses. Similar to the template evaluation, all the selected best models were subjected to further structural quality evaluation using the Ramachandran plot. All structural visualizations and analyses were carried out using Visual Molecular Dynamics (VMD) version 1.9.3 (Humphrey et Al., 1996), PyMOL by Schrödinger and Chimera (Pettersen et al., 2004). For predicting multimeric structure, the lowest MOF-containing structures were submitted to web-based program GalaxyHomomer (Baek et al., 2017).

### Gel filtration chromatography

Size exclusion gel filtration chromatography was carried out using purified recombinant PE11 on an analytical Superdex^TM^ 75 column connected to AKTApurifier chromatography unit (GE HealthCare). Briefly, 50 µg of purified PE11 protein was loaded onto the column equilibrated with 20 mM Tris, 150 mM NaCl pH 8.0 buffer in which protein was dialysed earlier. Elution was monitored at 280 nm. Four molecular weight markers- BSA (66kDa), carbonic anhydrase (29 kDa), cytochrome C (12.4 kDa) and aprotinin (6.5 kDa) were also loaded onto the column under similar conditions and elution volumes of respective markers were used to prepare a standard curve and compared with that of PE11 to deduce the molecular size.

### Colony morphology and Congo Red staining

Bacteria were cultured in 7H9 media until reaching mid-log phase, with an initial inoculum of 5 × 10^7^ colony forming units (CFU)/ml (OD580 nm: 0.1). Colony morphology was assessed by plating serial dilutions of cultures onto 7H11(BD BBL^TM^, USA) plates supplemented with glycerol and OADC, followed by 3 weeks of incubation at 37°C. To investigate cell wall properties, Congo red dye accumulation was evaluated following a protocol similar to Singh et al. (2016). Both Msmeg or M.tb strains were grown in 7H9 broth containing Congo red (Sigma-Aldrich, USA) and Tween 80, followed by cell harvesting, washing, and resuspension in acetone. After shaking, cells were pelleted, and the concentration of Congo Red in the supernatant was determined spectrophotometrically at 488 nm (EL808, Bio-Tek, USA).

### Isolation of mouse peritoneal macrophages

The selected C57BL/6 mice were maintained at the animal house facility of the Centre for DNA Fingerprinting and Diagnostics (CDFD). Experiments were performed as per the guidelines of the Institutional Animal Ethics Committee of CDFD. About 6 - 8 weeks old C57BL/6 mice were given one ml intraperitoneal injection of 4% thioglycolate medium. Three days post-injection, macrophages were harvested by peritoneal lavage as described earlier (Srivastava et al., 2019).

### Intracellular bacterial viability assay

The C57BL/6 peritoneal macrophages were infected with H37Rv, H37Rv-PE11KO, H37Rv- PE11KO::PE11, and H37Rv-PE11KO::PE11(S3) for 4 hours at an MOI of 1:10. post-infection, macrophages were washed with PBS containing Gentamycin at 50 µg/ml. The macrophages were then lysed in 0.1% TritonX at various time points, and serial dilutions of lysates were plated on 7H11 plates supplemented with 10% oleic acid, albumin, dextrose, catalase, and NaCl (HiMedia Laboratories). Plates were incubated at 37°C, and colonies were counted after 3–4 weeks (Srivastav et al., 2019).

## Results

### Biochemical characterization of rPE11

To check the biochemical activity of PE11/LipX protein, PE11 was overexpressed as a His-tagged protein in *E. coli* and isolated using Ni-NTA based affinity chromatography. The purified rPE11 protein showed a size of ∼11 kDa on a denaturing based gel electrophoresis (SDS-PAGE) similar to its predicted size based on amino acid sequence **(Figure 1A)**. PE11 has been previously reported to catalyze shorter and medium chain substrates ranging from C2 to C14 carbon but with highest preference for shorter acyl chain substrates C2-C4 (Singh et al., 2016). To assess its activity, we first determined the optimal pH and temperature conditions of recombinantly purified PE11 which were pH 8.0 and 37°C respectively using pNP-butyrate as substrate **(Supplementary Figures S1 A and B)**. Kinetic parameters of recombinant PE11 were measured by collecting time-dependent optical density data at various substrate concentrations using both pNP-acetate (C2) as well as pNP-butyrate (C4), and these values were used to construct a velocity vs substrate plot **(Figure 1B and 1C)**.

**Figure 1.**
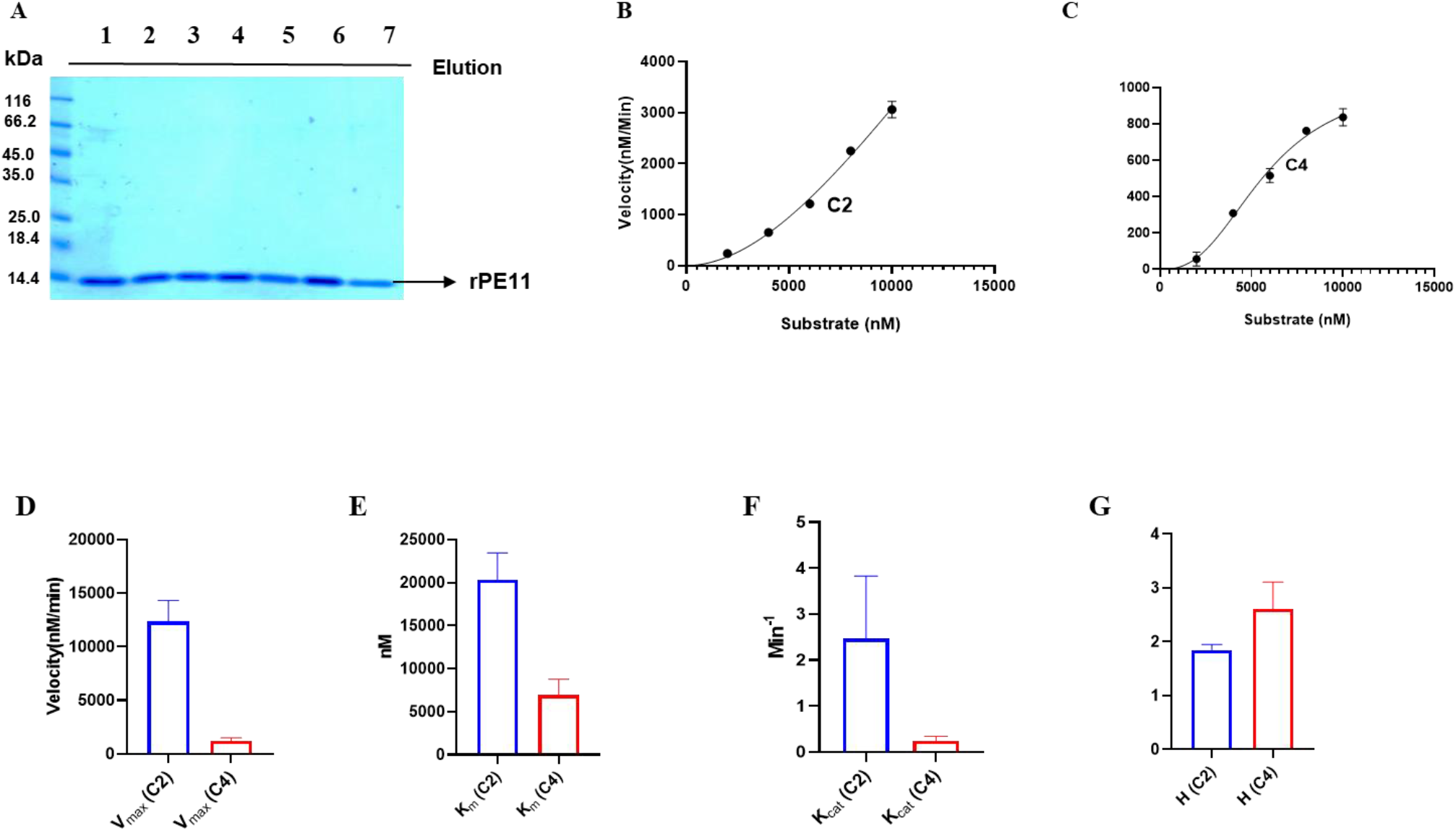
PE11 (LipX) is allosteric and shows positive co-operativity for binding substrates **(A)** Purification of recombinant His-tagged PE11 protein using NTA beads, shown on SDS-PAGE gel; Velocity vs Substrate graph for (**B**) pNP-acetate (C2) and (**C**) pNP-butyrate (C4) of recombinant PE11. **(D-G)** Comparative kinetics of recombinant PE11 for C2 and C4 substrate Vmax, Km, Kcat and H (Hill Coefficient). Data are representative of the mean ± SEM of three independent experiments. Statistical analysis was performed using one-way ANOVA. *P<0.05, **P<0.01 were considered significant.

The protein gives a sigmoidal curve with both substrates which served as the basis for calculating different kinetic parameters, including Km, Vmax, and Kcat using the GraphPad Prism© 9.1. Recombinant PE11 catalysed the hydrolysis of pNP-acetate with Vmax, Km, and Kcat values of 12334±3453 nM/min, 20362±5372 nM and 2.47±1.356 min^-1^, respectively and for pNP-butyrate those values were 1203±491.9 nM/min, 6929±319.3 and 0.24±0.98 respectively **(Figure 1D to 1F).** The mean Hill coefficient (H) values were 1.847 and 2.606 for pNP-acetate and pNP-butyrate respectively **(Figure 1G)**. Hill coefficient value of greater than 1 indicate that there is a positive cooperativity in substrate binding process. Furthermore, we examined the impact of different metal ions on the enzyme’s activity, focusing on both monovalent and divalent cations, namely potassium, calcium, manganese, and cobalt, with chloride ions acting as counter ions for all these salts. Our results demonstrated a significant increase in reaction velocity in the presence of monovalent potassium, while the enzyme exhibited reduced activity when exposed to other divalent metal ions **(Supplementary Figure S1C)**. Overall, these experiments suggest that PE11 is an esterase which catalyses hydrolysis of short chain esters (C2 and C4) and shows a clear preference for potassium as a metal ion that aids in its catalysis.

### PE11 protein shows signatures of both α-helix and β-sheets

Understanding the mechanism behind PE11’s mode of action requires insight into its protein structure. In the absence of any experimentally determined crystal structure of PE11 protein, we predicted the structure using ColabFold module based on AlphaFold (Mirdita et al., 2022). AlphaFold predicted a helix-loop-helix based structure with no beta strands (**Figure 2A**). To validate the AlphaFold-predicted secondary structural elements of PE11, the recombinantly purified protein was subjected to Circular Dichroism (CD) spectroscopy. CD analysis was performed in a phosphate buffer solution within a wavelength range of 260 nm to 190 nm at room temperature and pH 7.5. The obtained curve exhibited a discernible dip between 220 nm and 210 nm, indicative of the presence of α-helical structures (**Figure 2B**). Furthermore, a prominent positive crest was observed in the range of 200 nm to 190 nm, a characteristic of β-sheet structures. Importantly, this curve was consistent with typical characteristic features of the ‘αβ hydrolase’ fold family, featuring approximately ∼25% α-helix and ∼14% β-sheets, while the remaining structure comprised of random loops (∼30-%) and random structures (∼30%) based on analyses by JASCO **(Figure 2B-C)** (Ollis et al., 1992; Divya et al., 2018). Interestingly, this finding was in total contrast with that of AlphaFold based predictions which did not predict presence of β-sheet in PE11. Therefore, we reanalyzed the CD data using additional two independent programs (DichroWeb and BeStSel, an online program to analyze the CD data) as well as in different phosphate buffer systems. Surprisingly, all the programs indicated the presence of β-sheet structures ranging from 14-26% (**Figure 2C**). Since M.tb experiences a series of pH changes during its life cycle within the host, especially during phago-lysosomal fusion (Vandal et al., 2009), we speculated that AlphaFold predicted structure might represent the PE11 conformation for a specific pH. To address this, we analysed the PE11 protein at a pH range of pH 4.0 to pH 12.0 **(Figure 2D)**. Notably, within the range of pH 6.0 to pH 8.0, the obtained results converged into distinctive curves, signifying a stable structural configuration. Beyond this range, either lower or higher pH values led to structural disintegration/instability, resulting in the prevalence of random coils and sheets (**Figure 2D**). We subjected all these curves to Jasco, Dichroweb, and BeStSel programs to identify the secondary structural elements present at various pH. Unexpectedly, we could still detect the presence of beta sheets in all the pH range tested for the protein (**Figure 2E**). Overall, our CD-based biophysical experiments suggest the presence of both alpha helix and beta strands as distinct secondary structural elements in the PE11 protein.

**Figure 2.**
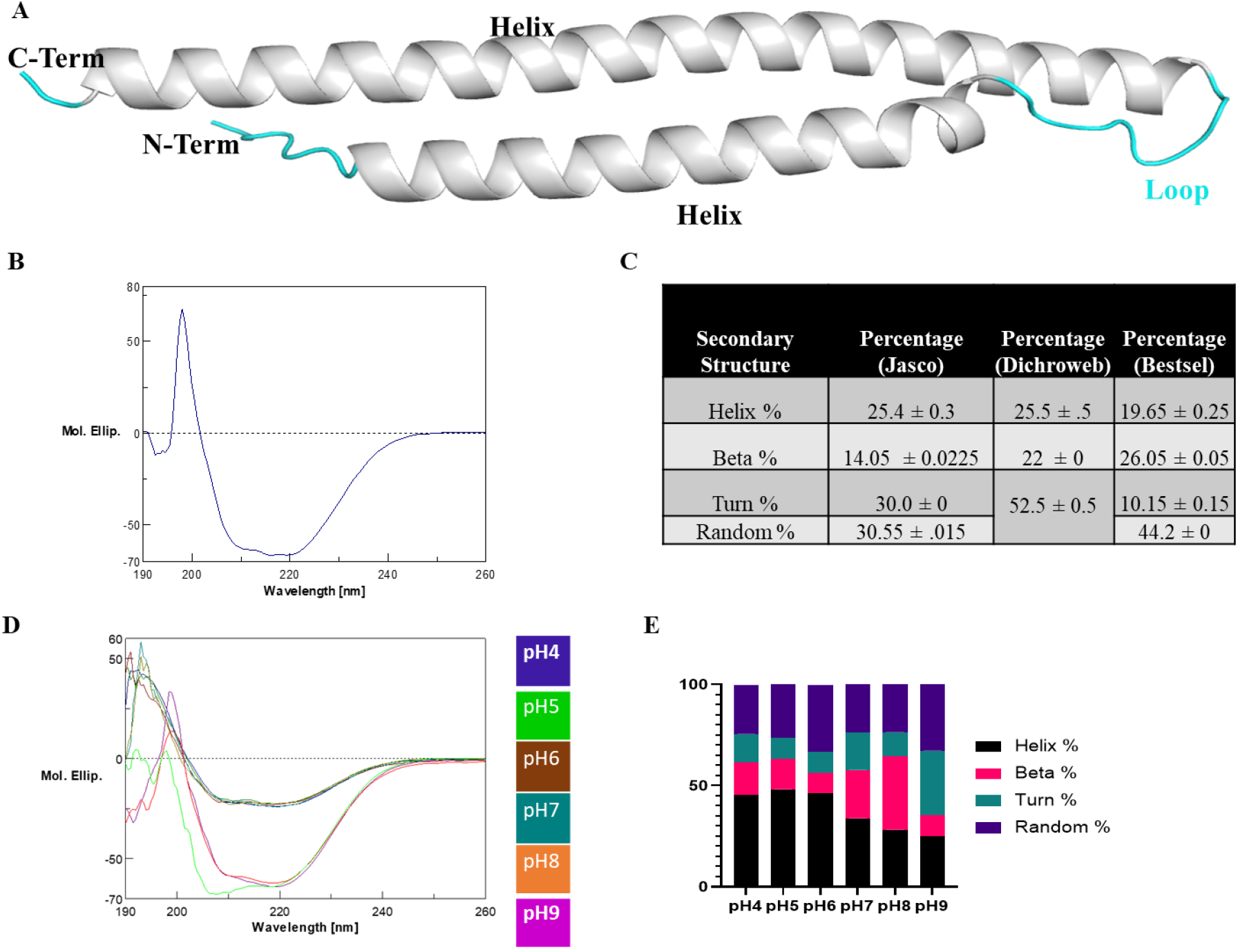
Recombinantly purified PE11 protein shows signatures of both α-helix and β- sheets (A) AlphaFold predicted structure of PE11 showing the presence of helix-turn-helix (HTH) structure. Structure was predicted using AlphaFold program via ColabFold (Mirdita et al., 2022). (**B**) CD spectra of recombinant PE11 protein in phosphate buffer at room temperature and pH 7.5. (**C**) Analysis of CD spectra through JASCO, DichroWeb, and BeStSel shows the presence of 20-25 α-helix and 14-25% β-sheets approximately. (**D**) CD spectra of recombinant PE11 protein at room temperature and a pH range of pH 4.0 to pH 9.0. (**E**) Graph showing the percentage of secondary structure elements calculated from CD spectra of PE11 at different pH. CD analysis was done with a fixed protein concentration of 0.30 mg/mL in phosphate buffer.

### Protein structure prediction using homology modelling

The disagreement of structural information of PE11 obtained from AlphaFold-based prediction and CD data is very intriguing. Therefore, a restraint-based homology modelling approach was adopted for developing all the theoretical structures and compared them using MODELLER© 10.0 (Webb and Sali, 2017; Webb and Sali, 2021). Earlier, Sultana et al. (2011) used a crystal structures of *Fusarium solani* cutinase (PDB_ID: 1CEX), MSMEG_6394, a lipase from *M. smegmatis* strain MC^2^155 (PDB_ID: 3AJA), a cutinase like protein from *Cryptococcus sp* (PDB_ID: 2CZQ) and mycobacteriophage esterase that cleaves the mycolylarabinogalactan bond to release free mycolic acids (PDB_ID: 3HC7). All the templates mentioned above were analyzed using a weighted pair group average clustering-based distance matrix, Modeller Objective Function (MOF) (Sánchez and Šali, 1997), Discrete Optimized Protein Energy (DOPE) (Shen and Sali, 2006), and GA341 scores (Eswar et al., 2008), and Ramachandran plot (Ramachandran and Sasisekharan, 1968) and were used for the multi template-based modelling. This resulted in the generation of five models for PE11. The lowest MOF-containing protein structure was selected for further analysis (**Figure 3A**). A Structural topology analysis **(Figure 3B and 3C)** suggested that the PE11 protein contained about 13% of beta-strand, 27% of alpha helices, and 58% of other structural formations **(Figure 3D)** which is in close agreement with the CD data analysed using JASCO earlier **(Figure 2C).** The predicted structure had 3 distinct β strands and 3 α-helices (**Figure 3A and 3B**) joined by loop like bends. The structural quality was in agreement with the Ramachandran plot. Overall, our structure prediction through MODELLER©10.0 and molecular dynamics-based simulation experiments are in agreement with our CD data, thereby suggesting a presence of α/β fold in PE11 protein.

**Figure 3.**
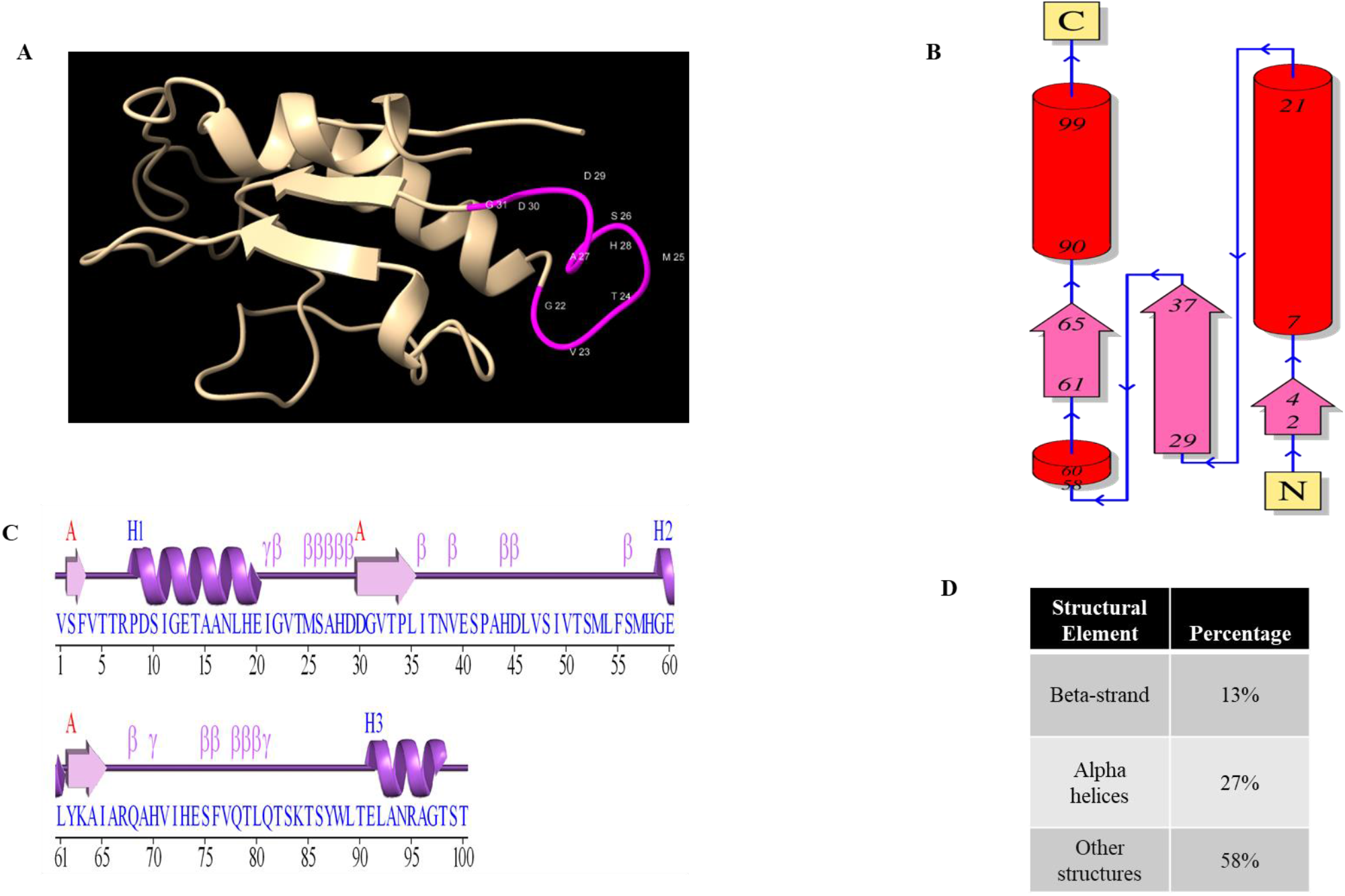
*In silico* template-based structure prediction of PE11 using MODELLER© 10.0 **(A)** Cartoon representation showing the lowest MOF-containing protein structure of the PE11 predicted using MODELLER© 10.0. Residues labeled in pink are the amino residues present (aa 22 to 31) in the homologous GX3SX4G motif containing the putative catalytic triad (**B**) Topology diagram showing the presence of different secondary structural elements of the lowest MOF-containing protein structure of the PE11 predicted using MODELLER© 10.0. (**C**) “Wiring diagram” of wild type PE11 protein showing distribution of α-helices and β-sheets (**D**) Distribution of structural elements in the lowest MOF-containing protein structure of the PE11 as predicted using MODELLER© 10.0.

### PE11 exist as dimer and tetramer

Earlier while determining the kinetics of PE11 enzymatic activity, we observed that Hill coefficient (H) values were more than 1 for both pNP-acetate and pNP-butyrate substrates. This indicated existence of positive cooperativity for binding of substrates to PE11, that is binding of one ligand facilitates binding of subsequent ligands at other sites of the enzyme. Given PE11 is small protein of ∼10.8 kDa, we dismissed the possibility multiple substrate binding sites on PE11. Therefore, we speculated existence of oligomeric forms of PE11 where binding of substrate in one chain may positively cooperate binding of substrate on other chains similar to binding of oxygen to haemoglobin (Ahmed et al., 2020). Accordingly, the purified protein was subjected to gel filtration chromatography using Superdex^TM^75 and

Tris-NaCl buffer. Two distinct elution peaks were observed, one corresponding to 40.6 kDa, indicating a tetrameric form of PE11 and another peak corresponding to 23.8 kDa, indicating a dimeric form of PE11 (**Figure 4A**). To further confirm our observations, we used web-based GalaxyHomomer program (Baek et al., 2017). *Ab initio* modelling in GalaxyHomomer indicated that PE11 possibly exist as dimer (**Figure 4B**), whereas template based-analysis indicated that PE11 can exist as a tetramer (**Figure 4C**).

**Figure 4:**
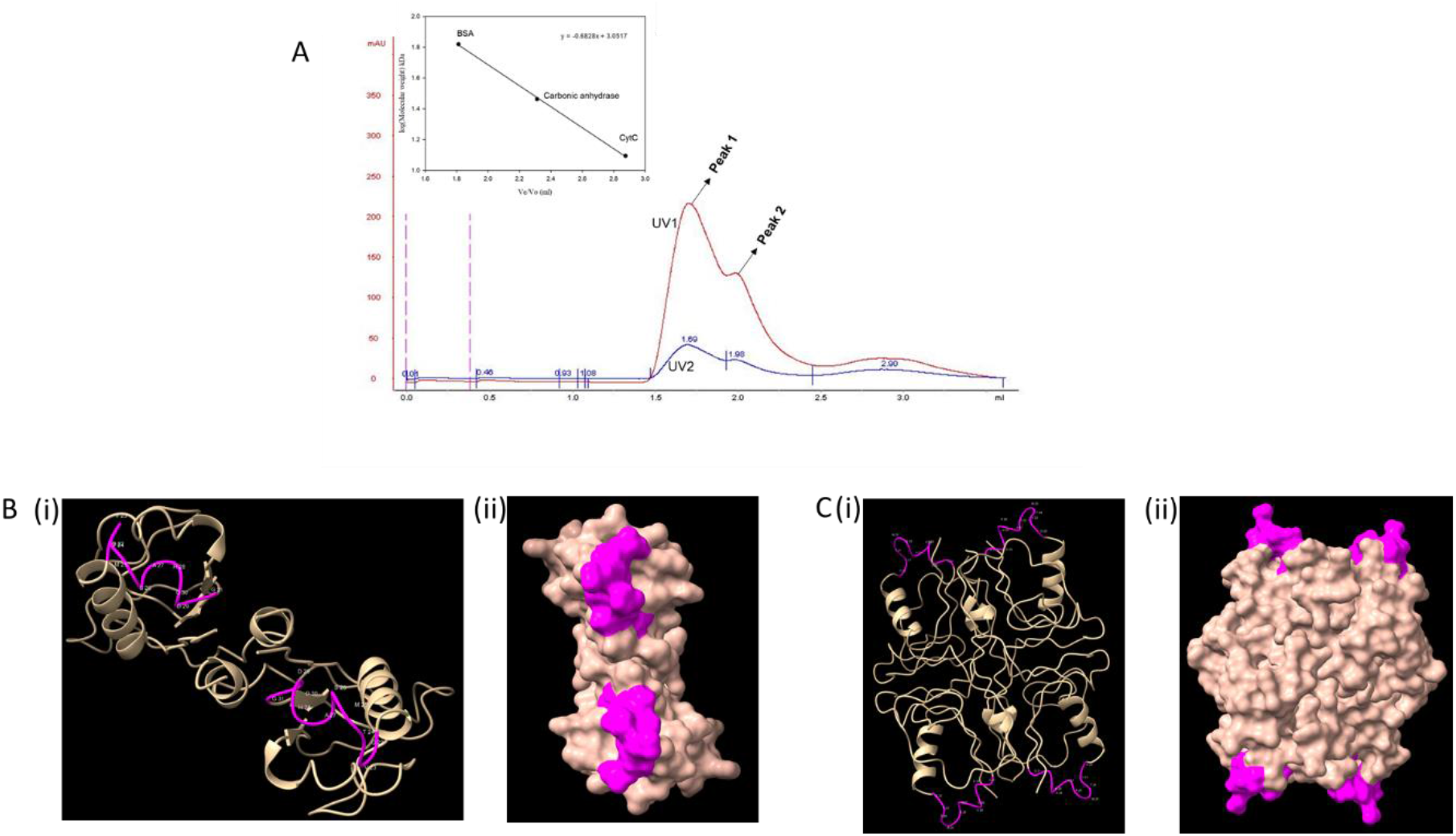
PE11 exists in oligomeric state (A) Size exclusion chromatography analysis of rPE11 revealed two distinct peaks, indicating presence of at least two oligomeric forms. The peaks correspond to molecular weights of 40.6 kDa (Peak 1) and 23.28 kDa (Peak 2), respectively, suggesting that PE11 exists in multiple oligomeric states. Inset, is the size standard calibration curve. (B) Cartoon showing dimeric conformation of PE11 as predicted by GalaxyHomomer where (i) Ribbon diagram (ii) Space filled diagram. (C) Cartoon showing predicted tetrameric conformation of PE11 as predicted by GalaxyHomomer where (i) ribbon diagram (ii) space filling model. Location of the amino acid residues encompassing the GX3SX4G motif is labelled in pink.

### S26 and G31 are crucial for PE11 esterase activity

Next, we aimed to identify the active site residues to understand the mechanism of esterase activity. Esterases are known to contain G-x-S-x-G motif (Bauer et al., 2019) typically present in many α/β hydrolase-fold family where the serine residue is part of the catalytic triad (Cao et al., 2015; Yang et al., 2019) in their active site. Interestingly, the sequence of the PE11 protein contained a homologous motif identified as G22X3S26X4G31 (**Figure 3A, highlighted in pink**) unlike typical GxSxG motif observed in serine hydrolases. To evaluate whether this motif is essential and part of the catalytic triad for esterase activity of PE11, we introduced substitution mutations in this motif where the Serine residue at 26^th^ position (S26) and the Glycine residue at 31^st^ position (G31) was replaced with Alanine [PE11(S3)] and examined its functional activity when expressed in *M. smegmatis.* When enzymatic assay was conducted using the whole cell lysates of all these bacilli using pNP-acetate as substrate, it was found that, PE11(S3) lost its esterase activity to the basal level similar to the control *M. smegmatis* carrying an empty vector **(Figure 5A).**

**Figure 5.**
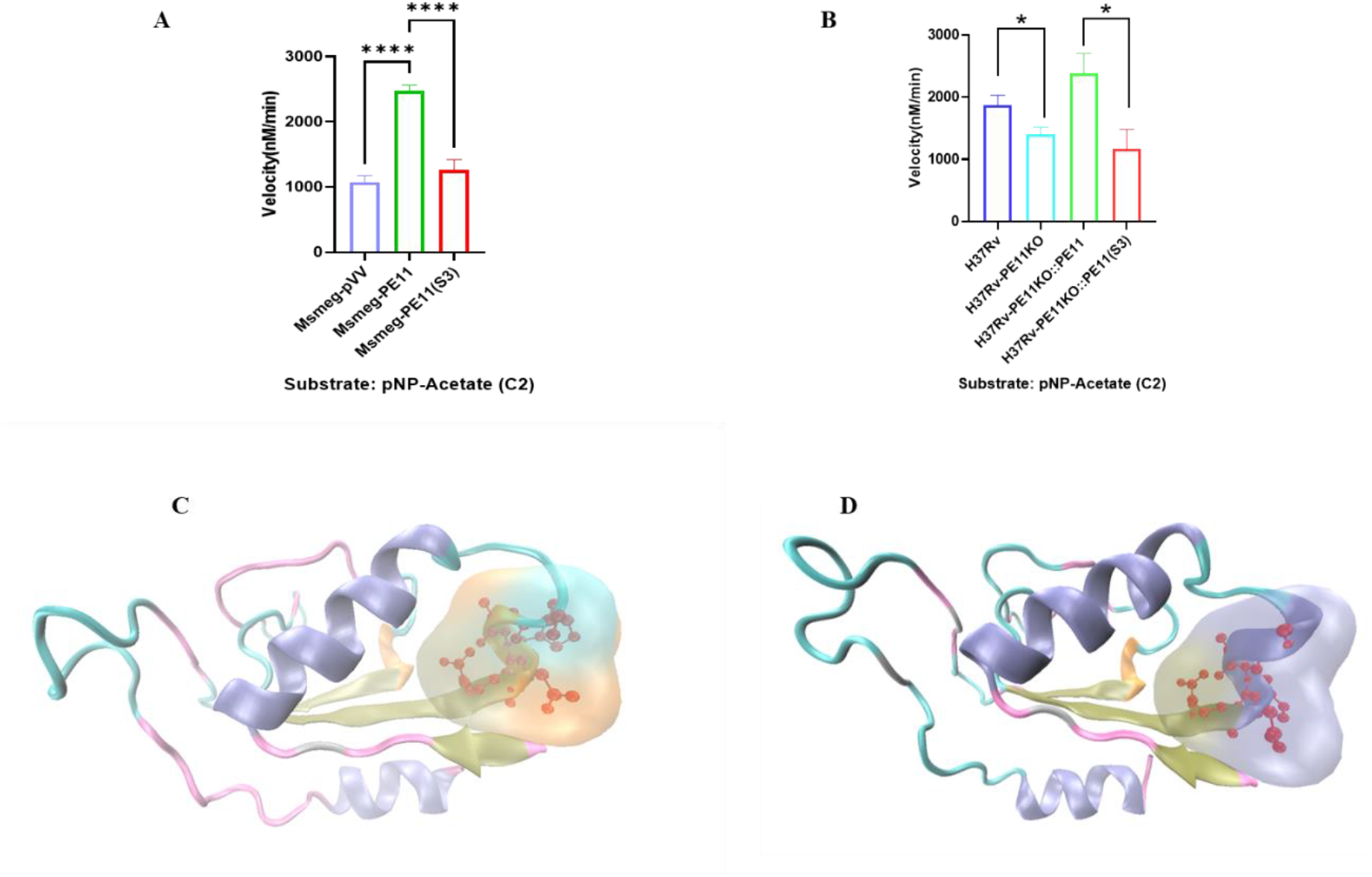
S26 and G31 residues are crucial for PE11 esterase activity (A) Enzymatic activity of *M. smegmatis* overexpressing wild-type PE11 and the mutant PE11(S3), in the form of pNP released per nM/min using respective *M. smegmatis* cell lysates for using pNP-acetate (C2) as substrate. (B) Enzymatic activity of cell lysates prepared from wildtype *M. tuberculosis* strain H37Rv, PE11 knock-out strain (H37Rv-PE11KO), H37Rv- PE11KO complemented with wild-type PE11 (H37Rv-PE11KO::PE11) and H37Rv-PE11KO complemented with mutant PE11(S3) [H37Rv-PE11KO::PE11(S3)] in the form of pNP released per nM/min using pNP-acetate (C2) as substrate. Data are representative of the mean ± SEM of 3 different experiments. Statistical analysis was performed using one-way ANOVA. *P<0.05, **P<0.01 were considered significant. (C) Cartoon representing space filling model of the putative catalytic triad region of wild type PE11 and (D) PE11(S3) mutant.

To further support these observations, we generated PE11 knocked out mutants using *M. tuberculosis* H37Rv strain (H37Rv-PE11KO) and strains complemented with either the wild-type PE11 or mutant PE11(S3) (**Supplementary Figure S2**) and carried out esterase assay using pNP-acetate as substrate. The results indicated that H37Rv-PE11KO complemented with mutant PE11(S3) [H37Rv-PE11KO::PE11(S3)] also lost activities to the basal level of H37Rv-PE11KO whereas the wild-type H37Rv and H37Rv-PE11KO complemented with wild-type PE11 (H37Rv-PE11KO::PE11) had higher and comparable enzymatic activities (**Figure 5B).** Thus, these findings indicate that the S26 and G31 residues are critical for the esterase activity of PE11 and possibly form part of the catalytic triad.

For several hydrolases, the catalytic triad containing the Serine residue (S) from GxSxG motif along with Aspartic acid (D) and Histidine (H) is known to constitute the active catalytic site. It was speculated that S26 along with H28 and D29/D30 are likely to be the part of the putative catalytic triad which are present within the G22X3S26X4G31 motif due to their proximity and sequential orientation **(Figure 3A)** (Sultana et al., 2011; Yang et al., 2019). To further investigate impact of amino acid substitution in the putative catalytic triad, we determined the 3D structure of the mutant PE11(S3) as described earlier. It was found that, S26A and G31A substitutions brought steric conformational changes in the catalytic triad pocket of PE11 which explains possible loss of enzymatic function in mutant PE11 (compare **Figures 5C** **with 5D**).

### PE11 esterase activity enhances resistance to antibiotics

The mycobacterial cell envelope boasts a formidable, lipid-rich, hydrophobic coat. The unique structural feature of the cell wall not only provides an efficient shield against antibiotics (Daffe and Draper, 1998), but also mycobacteria carry out vital functions such as safeguarding the bacterial cell against hostile environmental conditions such as in host phagosome after infection (Harding and Boom, 2010; Murry et al., 2009; Reed et al., 2004). Moreover, it plays a pivotal role in defining crucial virulence attributes of this pathogen. *M. smegmatis* expressing PE11 (Msmeg-PE11) showed higher persistence against various stressors such as SDS, H2O2, lysozyme and various antibiotics (Singh et al., 2016). Therefore, we further investigated whether esterase activity of PE11 is actually responsible for higher persistence of the bacterium against antibiotics. Accordingly, the esterase mutant Msmeg- PE11(S3) was cultured in the presence of Rifampicin and Isoniazid (3:2 ratio, 20 µg/ml) and compared its rate of survival with the wild-type control (Msmeg-PE11) and *M. smegmatis* harbouring an empty vector (Msmeg-pVV). Expectedly, the wild-type Msmeg-PE11 showed high persistence whereas mutant Msmeg-PE11(S3) with lower enzymatic failed to grow in the presence of antibiotics suggesting a role of esterase activity in providing resistance against various stressors **(Figure 6A)**.

**Figure 6.**
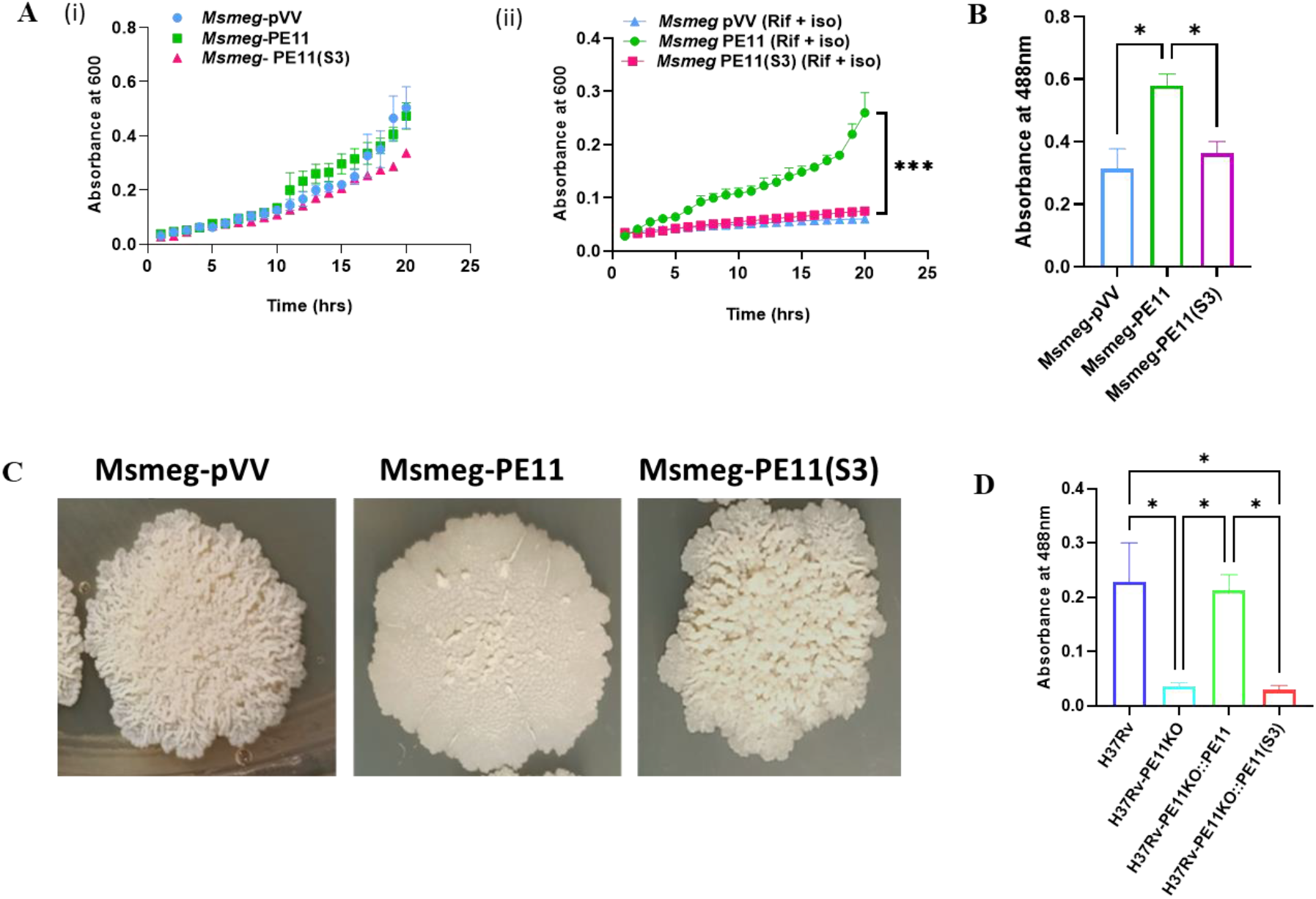
PE11 esterase activity provide Antibiotic resistance and alters cell wall architecture **(A)** Growth of Msmeg-pVV, Msmeg-PE11, and Msmeg-PE11(S3) bacilli in Absence **(i)** or presence **(ii)** of Rifampicin and Isoniazid (3:2, 20 µg/ml) was analyzed for 24 hours, **(B)** In a separate experiment, cultures of Msmeg-pVV, Msmeg-PE11, and Msmeg-PE11(S3) were grown in 7H9 broth containing 100 μg/ml of Congo Red and 0.05% Tween 80 at 37°C for 24 hours. The concentration of Congo Red in the supernatant after incubating the pellet in acetone was quantified using spectrophotometry at 488 nm. **(D)** Cultures of H37Rv, H37Rv- PE11KO, H37Rv-PE11KO::PE11 and H37Rv-PE11 KO::PE11(S3) were grown in 7H9 broth containing 100 μg/ml of Congo Red and 0.05% Tween 80 at 37°C for 24 hours. Pellets were resuspended in acetone for 2 hours and the concentration of Congo Red in the supernatant was quantified using spectrophotometry at 488 nm. Data are representative of the mean ± SEM of different experiments. Statistical analysis was performed using one-way ANOVA. *P<0.05, **P<0.01 were considered significant.

### PE11 esterase activity alters cell wall architecture and colony morphology of mycobacteria

PE11 is known to enhance hydrophobicity of the cell envelop when overexpressed in *M. smegmatis* (Singh et al., 2016). To further confirm the role of PE11 esterase domain in modulating lipid content, cell wall composition and hydrophobicity, we used Congo Red (a hydrophobic diazo-dye known for binding to lipids and lipoproteins). It was found that, Msmeg-PE11(S3) had reduced accumulation of Congo Red due to decreased hydrophobicity (**Figure 6B**). As there is a link between increased lipid content and altered cell wall structure influencing colony morphology (Singh et al., 2016), we next examined the colony morphology of the bacilli to gain further insights into the morphological effects of S26A and G31A substitution in PE11. Colonies were plated, incubated at 37°C, and photographed after 4-5 days. Strikingly, the distinctive colony morphology observed in Msmeg-PE11-positive transformants was lost in Msmeg-PE11(S3). While colonies of Msmeg-pVV exhibited typical irregular and rough structures, those of Msmeg-PE11 were rounded and smoother. Conversely, Msmeg-PE11(S3) colonies closely resembled those of Msmeg-pVV, rather than Msmeg-PE11 (**Figure 6C**).

M.tb exhibits a complex array of enzymes engaged in similar metabolic functions, yet the question of their redundancy or distinct roles remains a focal point to understand their specific roles in mycobacterial pathology. To investigate the specific role of PE11 esterase activity in the regulation of M.tb cell wall architecture, Congo Red accumulation assay was carried out with H37Rv, H37Rv-PE11KO, and complemented strains. The results indicated higher accumulation of Congo Red in H37Rv and H37Rv-PE11KO::PE11, as compared to that in H37Rv-PE11KO and H37Rv-PE11KO:: PE11(S3) (**Figure 6D**). These data indicate that PE11 esterase activity plays a crucial role in M.tb cell wall remodelling.

### PE11 esterase activity provides a survival advantage to both *M. smegmatis* and *M. tuberculosis*

Modulation of cell wall architecture can influence the virulence efficiency of mycobacteria. Therefore, we next investigated the impact of PE11 esterase activity on mycobacterial survival inside macrophages. Thus, peritoneal macrophages from C57BL/6 mice were infected with Msmeg-pVV or Msmeg-PE11 or Msmeg-PE11(S3) at an MOI of 1:10 for 3, 24, and 48 hours. This was aimed to assess the relationship between enzymatic activity and virulence, as well as bacterial viability post-infection. CFU counts indicated better survival of Msmeg-PE11 as compared to both Msmeg-PE11(S3) and Msmeg-pVV, suggesting the vital role of PE11 esterase activity in pathogen survival within macrophages (**Figure 7A**).

**Figure 7.**
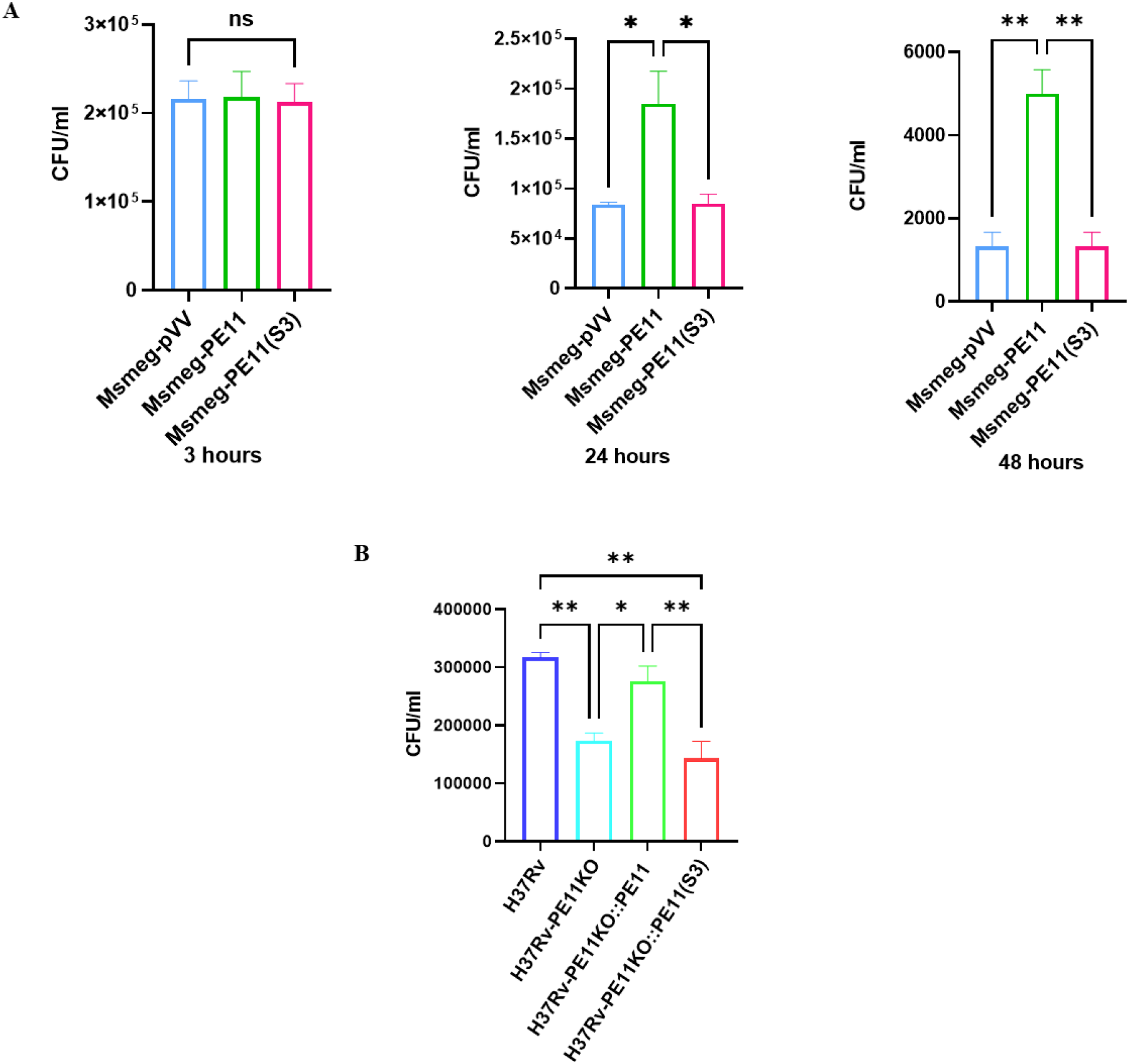
PE11 esterase activity provides survival advantage to Mycobacteria. C57BL/6 mice peritoneal macrophages were infected with either **(A)** Msmeg-pVV/Msmeg- PE11/Msmeg-PE11(S3) at 1:10 M.O.I for 3, 24 and 48 hours or **(B)** H37Rv/H37Rv- PE11KO/H37Rv-PE11KO::PE11/H37Rv-PE11 KO::PE11(S3) at 1:10 M.O.I for 24 hours. Following infection, the cells were lysed and CFU counts were monitored. Data are representative of the mean ± SEM of different experiments. Statistical analysis was performed using one-way ANOVA. *P<0.05, **P<0.01 were considered significant.

Similar experiment was carried out using M.tb PE11 knockout (H37Rv-PE11KO) and complemented strains in order to assess the role of PE11 enzymatic activity in M.tb virulence. Accordingly, peritoneal macrophages were infected with H37Rv, H37Rv-PE11KO, H37Rv-PE11KO::PE11, and H37Rv-PE11KO::PE11(S3) at 1:10 MOI. A similar assessment of intracellular survival revealed higher CFU counts in H37Rv and H37Rv-PE11KO::PE11 compared to H37Rv-PE11KO and H37Rv-PE11KO::PE11(S3), indicating that PE11 esterase activity confers a survival advantage to the bacilli (**Figure 7B**).

## Discussion

The proline-glutamic acid (PE)/proline-proline-glutamic acid (PPE) proteins play a vital role in *M. tuberculosis* virulence (Sampson, 2011). Investigating esterases/lipases, particularly from the PE family, is particularly challenging due to difficulties in obtaining enzymatically active forms via the *E. coli* expression system, often resulting in inactive inclusion bodies. Overcoming this requires effective renaturation (Garrett et al., 2015; Sultana et al., 2013)

Lipids and free fatty acids are important sources of energy for *M. tuberculosis* during infection and latency and are significantly involved in the virulence (Lin et al., 2024). Out of the 24 members of the Lipase (Lip) family, few of them have been shown to exhibit esterase activity on shorter carbon chain substrates (Lin et al., 2024). The metabolic utilization of SCFA (Small chain fatty acids) appears to be beneficial factor for the improved growth of *Mycobacterium* sp., emphasizing the supportive role of SCFA-metabolizing esterases in enhancing mycobacterial persistence (Silva et al., 2020). Among the Lip family of proteins of *M. tuberculosis*, the role of PE11 in lipid synthesis was emphasized by a study by Rastogi et. al (2017) where PE11 knock-down significantly reduced PDIM levels, implicating PE11 to be involved directly in PDIM synthesis. PDIMS are crucial for *M. tuberculosis*’s ability to escape phagosomes, induce host cell necrosis, and trigger macro autophagy, enhancing its virulence (Quigley et al., 2017). Earlier, we showed that PE11 has esterase activity which is essential for cell wall remodelling and preferentially hydrolyses short chain esters (Singh et al., 2016), however, detailed biochemical and structural characterisation was not carried out earlier.

When we compared short chain esters pNP-acetate (C2) and pNP-butyrate (C4) as its substrate, we found that Vmax and Kcat is significantly higher for C2 esters as compared to C4 esters but affinity for C4 ester is more as indicated by lower Km values indicating that PE11 can utilize C4 at lower available substrate concentrations. We also found that Vmax increases when K^+^ ions are present. Interestingly, intraphagosomally K^+^ concentrations surge during the maturation of macrophage phagosomes (MacGilvary et al., 2019). M.tb experiences different pH during its growth conditions. Therefore, a broad pH spectrum (pH 4.0–12.0) was used to determine enzyme activities. At extreme pH levels (pH 4.0 and pH 9.0), minimal activity was observed, likely due to partial or complete denaturation of PE11 enzyme, particularly in its alpha-helical structure. This suggests that PE11 activity relies on its conformation. Similar observations were made for Rv0045c, LipL, LipE, LipC and other mycobacterial esterase/lipases (Shen et al., 2012, Cao et al. 2015, Yang et al., 2019).

Among the lipase/esterase family of proteins of *M. tuberculosis*, PE11 (LipX), is the smallest protein of molecular mass about 10.8 kDa and lacks the canonical “GxSxG” motif (Deb et al., 2006). The conventional approach of template selection through BLAST and PSI-BLAST was able to select templates with 2 large alpha helices and a loop in between **(Figure 2A).** However, subsequent analysis using CD spectra and insights from protein family information indicated presence of beta sheets within the protein’s structural architecture. As a result, we chose to employ homology modeling approach using templates as described by Sultana et al., (2011), to predict the structure of PE11. The resulting model revealed a composite structure comprising α-helices, β-sheets, β-turns, and other elements. The PE11 protein, categorized within the α/β hydrolase fold family according to SCOP (Murzin et al., 1995), displays a structural arrangement at the α/β/α level **(Figure 3A).** However, the model lacked typical “lid” like structure that controls the access to the active site (Kumar et al., 2017). Being a small protein, we expected less elements of structural complexity to be present in PE11. Since, PE11 did not contain a conserved canonical G-x-S-x-G motif characteristic of several esterases/lipases (Bauer et al., 2019), we predicted a homologous GX3SX4G motif encompassing amino acid residues from 22-31.

Typically, the serine residue (S) from GxSxG motif along with aspartic acid (D) and a Histidine residue (H) is known to constitute the active catalytic triad (Sultana et al., 2011; Yang et al., 2019). Based on the presence of Serine^26^ in a bend between α-helix (aa residues 7-21) and a β-strand (aa residues 29-37) (**Figure 3A and 3C**) (Kumar et al., 2017), we predicted S26 residue to be part of the active catalytic triad. We also predicted that H28 and D29/D30 are likely to be part of the putative catalytic triad which are present within the GX3SX4G motif due to their proximity (**Figure 3A**). However, the sequential orientation of the catalytic triad appears to be ser-his-asp rather than ser-asp-his as found in many mycobacterial esterase/lipases (Sultana et al., 2011; Li et al., 2017; Yang et al., 2019). Spatially distant catalytic triad residues are likely to accommodate long chain substrates whereas adjacent residues are likely to prefer short-chain substrates (Li et al., 2017). Since, PE11 was found to prefer short-chain (C2) substrates, we speculated that S26, H28 and D29/30 are part of the catalytic triad as other Histidine and Aspartic acid residues were found to be located in a larger distance spatially.

In order to validate our hypothesis, we substituted S26 and G31 residues with neutral Alanine residue [PE11(S3)] and compared its activity with the wild-type PE11. It was found that the mutant PE11(S3) lost almost all of its enzymatic activity was observed in whole cell lysate **(Figure 5A).** Similarly, when a H37Rv-PE11KO strain was complemented with wild-type PE11 (H37Rv-PE11KO::PE11), enzymatic activity was comparable with that of wild-type H37Rv, but enzymatic activity was found to be reduced to the level of H37Rv-PE11KO when the same strain was complemented with PE11(S3) [H37Rv-PE11KO::PE11(S3)] (**Figure 5B**). Homology modelling of PE11(S3) showed that, there is a significant conformational change around the putative catalytic triad due to Alanine substitution at S26 and G31, which explains the loss of enzymatic activity of PE11(S3) (**Figures 5C and 5D**). These observations highlight the functional significance of the predicted GX3SX4G motif and location of the putative catalytic triad within this motif.

Interestingly, while analyzing the enzyme kinetics data we found that the Hill Coefficient (H) values were > 1 indicating positive cooperativity of binding of substrates which is characteristic of allosteric enzyme having multiple binding sites for substrates/ligands. Since PE11 is a small protein of about 100 amino acids (∼10.8 kDa), and we found only one putative catalytic triad of ‘GX3SX4G’, we speculated could be multimeric enzyme to show such positive cooperativity of binding. Accordingly, we performed gel filtration chromatography and we found two major peaks corresponding to the sizes of 40.6 kDa and 23.28 kDa indicating that PE11 can exist as tetramer and dimer. When the lowest MOF structure of PE11 was analysed using GalaxyHomomer, it predicted dimeric organization of PE11 using *ab initio* docking without using any template information. In the highest-score structure returned by the program shows that the C-terminal α-helices interact with each other to form the dimer (**Figure 4B**). On the other hand, the template-based prediction module, returned a tetrameric conformation of PE11 enzyme (**Figure 4C**). Therefore, these data indicated that PE11 exist as dimer and tetramer, which explains the observed H value of >1.

Furthermore, the physiological implications of PE11 esterase activity as well as the predicted critical amino acid residues in mycobacterial bacilli was studied. We observed the loss of LipX characteristic cell wall architecture, and lipid profile in the PE11(S3) mutant strain suggesting the crucial role of predicted amino acids in the active site of the protein. The mycobacterial cell envelope is lipid-rich and acts as a hydrophobic shield and thereby is able to provide defence against antibiotics and environmental challenges. As emphasized by Alderwick et al. (2015), this resilient structure plays a critical role in preserving bacterial integrity. *M. tuberculosis* possesses a distinctive cell wall architecture, featuring mycolic acid, trehalose dimycolate, PDIMs, PGL, and cord factor, which collectively form a hydrophobic barrier. This barrier hampers antibiotic diffusion, contributing significantly to drug resistance (Ghazaei, 2018). In conditions of nutrient limitation, the production of the outer membrane lipid phthiocerol dimycocerosate (PDIM) becomes indispensable for antibiotic tolerance. Mutants exhibiting altered antibiotic tolerance, identified through a transposon mutant library, highlight the crucial role of PDIM as a key determinant in M.tb drug tolerance (Block et al., 2023). Previous studies have associated PE11 knock down with a decrease in PDIMs in M.tb and enhanced persistence by *M smegmatis* over-expressing PE11 against various stressors and antibiotics (Rastogi et al., 2017; Singh et al., 2016). Our results reiterated these findings as PE11(S3) expressing *M. smegmatis* had higher sensitivity to antibiotics and altered colony morphology. Notably, even in the presence of combination drugs like rifampicin and isoniazid, the research highlights the critical role of PE11/LipX esterase activity in antibiotic tolerance, further validating the potential of predicted sites for therapeutic interventions.

M.tb often has multiple enzymes performing similar metabolic tasks, yet it remains unclear if these enzymes are interchangeable or have distinct functions. The question arises whether there is functional redundancy contributes to specific aspects of the cellular process. The essentiality of the PE11 gene has been a topic of debate, with Griffin et al. (2011) and DeJesus et al. (2017) presenting differing assertions in their studies. Nonetheless, in our present study, a knockout of the PE11 gene in M.tb was effectively generated, offering evidence that supports its non-essential role. Despite the protein’s evident role in the virulence and pathogenicity of the pathogen, our findings indicate that PE11 is critical but not essential for bacterial survival in culture.

Lipases that respond to stress, such as LipD (reacting to oxidative stress), LipU (responsive to nutritive stress), and LipY (dormancy and lipid metabolism), play pivotal roles in navigating challenging environments encountered during infection in the host milieu (Li et al., 2017; Singh et al., 2014; Santucci et al., 2019). An observable difference can be noted in the colonies of Msmeg-bacilli having overexpressed PE11 protein when compared to Msmeg- pVVas well as Msmeg-PE11(S3). This difference hints towards involvement of PE11 esterase activity in cell wall-related metabolism. Furthermore, peritoneal macrophage infection-based bacterial viability study revealed that PE11 provides a distinct survival advantage to both M.tb and Msmeg through its esterase activity.

These comprehensive findings emphasize the significance of PE11 esterase activity as well as the predicted sequence on bacterial morphology, hydrophobicity, and virulence in both *M. smegmatis* and *M. tuberculosis*. This insight provides valuable perspectives into mycobacterial physiology, unveiling potential avenues for therapeutic interventions. Notably, the predicted catalytic site of PE11 emerges as a promising target for small molecule inhibitors in combination with primary drugs or alone to enhance the present treatment for tuberculosis. Our present study was successful in characterizing and deciphering the sequence and structural components PE11 protein of *M. tuberculosis*.

## Acknowledgement

The authors thank Dr. Madhu Battu and Dr. Manoj Bisht for help. The authors gratefully acknowledge the financial support by the Science and Engineering Research Board (SERB), Department of Science and Technology (DST), Government of India (JCB/2021/000035), Indian Council of Medical Research (ICMR), Govt. of India (2021-10087/GTGE/ADHOC-BMS) and a core grant from CDFD by DBT. PD is supported by the fellowship from Council of Scientific and Industrial Research (CSIR), Govt. of India.

---

**Supplementary Figure S1.**
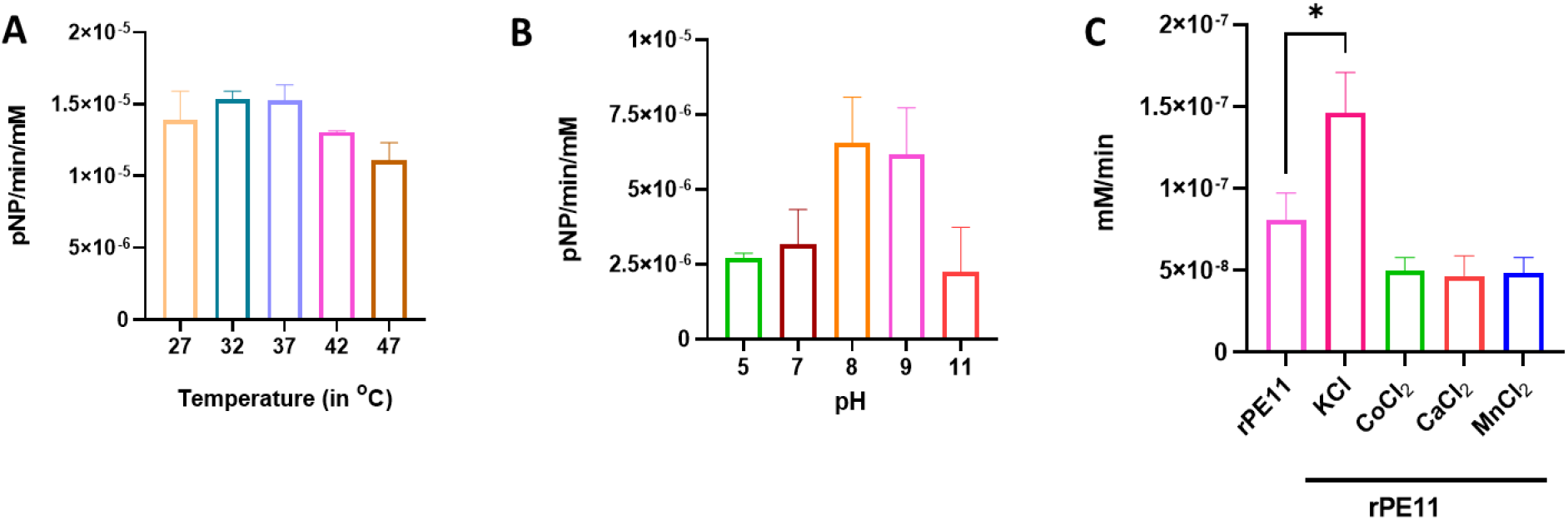
Biochemical characterization of the recombinant PE11 (LipX) protein Effect of (A) temperature and (B) pH (C) different metal ion (5 µM) on the esterase activity of PE11. Data are representative of the mean ± SEM of different experiments. Statistical analysis was performed using one-way ANOVA. *P<0.05, **P<0.01 were considered significant.

**Supplementary Figure S2:**
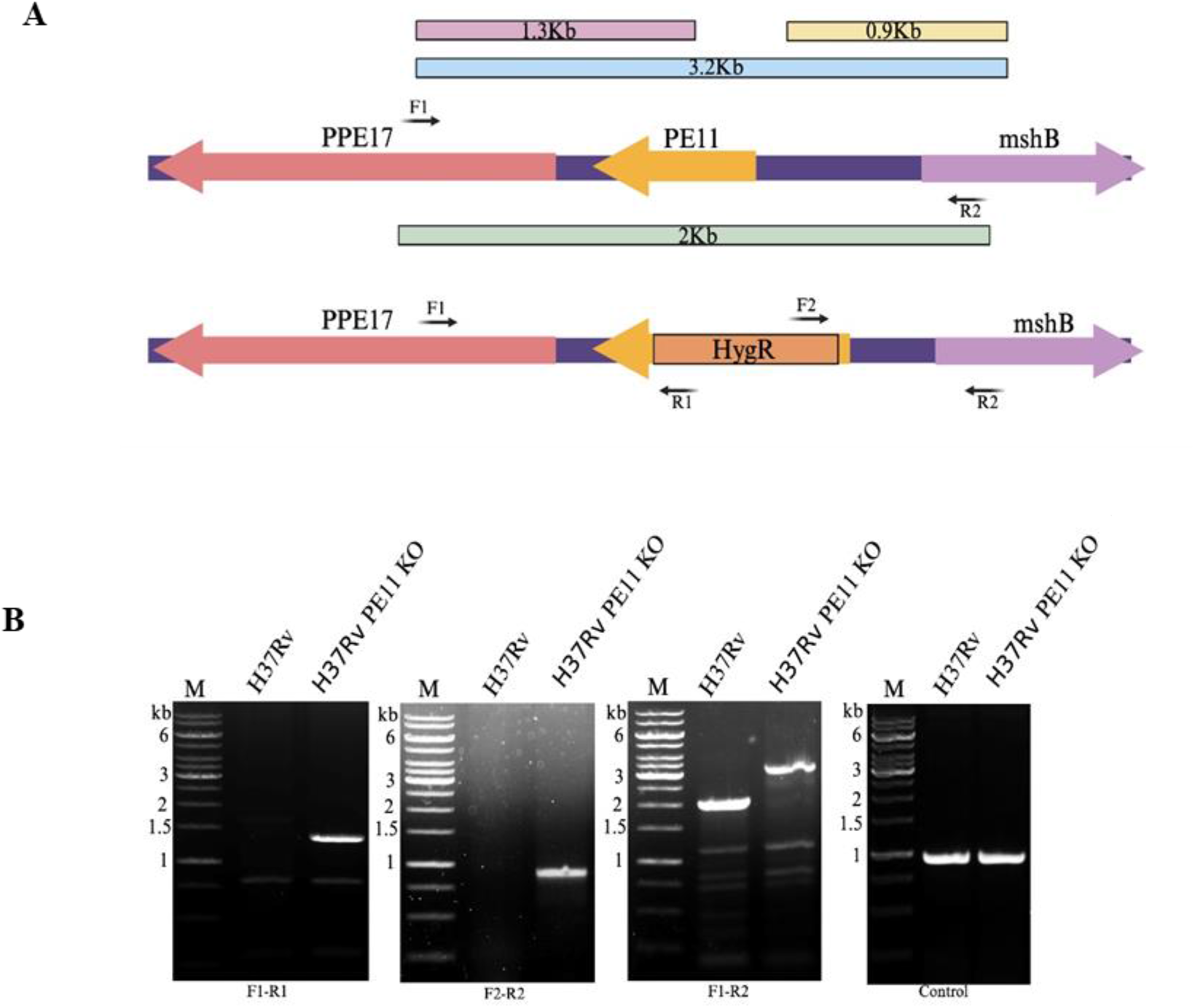
Generation of PE11 Knocked Out (PE11-KO) H37Rv strain of *M. tuberculosis* **(A)** Schematic representation of location and domain architecture of PE11 gene in *M. tuberculosis*, along with depiction of the homologous recombination process between the Allelic Exchange Substrate (AES) and the native genomic loci of PE11 (Rv1169c). **(B)** Confirmation of authenticity of recombination by PCR assays, using the depicted primers for validation, particularly F1-R1, F2-R2, and F1-R2 which shows a notable shift in the band size from 1.2 kb to 2 kb after the replacement of PE11 gene by Hygromycin resistance gene (Hyg^R^).

## References

1. Ahmed MH, Ghatge MS, Safo MK. Hemoglobin: Structure, Function and Allostery. Subcell Biochem. 2020;94:345–382.

2. Alderwick, L. J., Harrison, J., Lloyd, G. S., & Birch, H. L. (2015). The Mycobacterial Cell Wall--Peptidoglycan and Arabinogalactan. Cold Spring Harbor Perspectives in Medicine, 5(8), a021113.

3. Altschul, S. F., Madden, T. L., Schäffer, A. A., Zhang, J., Zhang, Z., Miller, W., … & Lipman, D. J. (1997). Gapped BLAST and PSI-BLAST: a new generation of protein database search programs. Nucleic Acids Research, 25(17), 3389–3402.

4. Baek M, Park T, Heo L, Park C, Seok C. GalaxyHomomer: a web server for protein homo-oligomer structure prediction from a monomer sequence or structure. Nucleic Acids Res. 2017 Jul 3;45(W1):W320–W324.

5. Bauer, T. L., Buchholz, P. C. F., & Pleiss, J. (2020). The modular structure of α/β- hydrolases. FEBS Journal, 287(5), 1035–1053.

6. Betts, J. C., Lukey, P. T., Robb, L. C., McAdam, R. A., & Duncan, K. (2002). Evaluation of a nutrient starvation model of *Mycobacterium tuberculosis* persistence by gene and protein expression profiling. Molecular Microbiology, 43(3), 717–731.

7. Block, A. M., Namugenyi, S. B., Palani, N. P., Brokaw, A. M., Zhang, L., Beckman, K. B., & Tischler, A. D. (2023). *Mycobacterium tuberculosis* Requires the Outer Membrane Lipid Phthiocerol Dimycocerosate for Starvation-Induced Antibiotic Tolerance. mSystems, 8(1), e0069922.

8. Brennan, M. J., & Delogu, G. (2002). The PE multigene family: a ’molecular mantra’ for mycobacteria. Trends in Microbiology, 10(5), 246–249.

9. Cao, J., Dang, G., Li, H., Li, T., Yue, Z., Li, N., … & Chen, L. (2015). Identification and Characterization of Lipase Activity and Immunogenicity of LipL from *Mycobacterium tuberculosis*. PLoS One, 10(9), e0138151.

10. Cole, S., Brosch, R., Parkhill, J., et al. (1998). Deciphering the biology of *Mycobacterium tuberculosis* from the complete genome sequence. Nature, 393, 537–544.

11. Daffe, M., & Draper, P. (1998). The envelope layers of mycobacteria with reference to their pathogenicity. Advances in Microbial Physiology, 39, 131–203.

12. Deb C, Daniel J, Sirakova TD, Abomoelak B, Dubey VS, Kolattukudy PE. A novel lipase belonging to the hormone-sensitive lipase family induced under starvation to utilize stored triacylglycerol in Mycobacterium tuberculosis. J Biol Chem. 2006 Feb 17;281(7):3866–75.

13. DeJesus, M. A., Gerrick, E. R., Xu, W., Park, S. W., Long, J. E., Boutte, C. C., … & Ioerger, T. R. (2017). Comprehensive analysis of the *Mycobacterium tuberculosis* genome via saturating transposon mutagenesis. mBio, 8(1), e02133–16.

14. Delogu, G., & Brennan, M. J. (2001). Comparative immune response to PE and PE_PGRS antigens of *Mycobacterium tuberculosis*. Infection and Immunity, 69(9), 5606–5611.

15. Deng, W., Long, Q., Zeng, J., Li, P., Yang, W., Chen, X., & Xie, J. (2017). *Mycobacterium tuberculosis* PE-PGRS41 Enhances the intracellular survival of *M. smegmatis* within macrophages via blocking innate immunity and inhibition of host defense. Scientific Reports, 7, 1–13.

16. Divya, M. B., Vemula, M., Balakrishnan, K., Banerjee, S., & Guruprasad, L. (2018). *Mycobacterium tuberculosis* PE1 and PE2 proteins carrying conserved α/β-serine hydrolase domain are esterases hydrolyzing short to medium chain p-nitrophenyl esters. Progress in Biophysics and Molecular Biology, 140, 90–102.

17. Eswar, N., Eramian, D., Webb, B., Shen, M. Y., & Sali, A. (2008). Protein structure modeling with MODELLER. In Structural proteomics: high-throughput methods (pp. 145-159). Springer.

18. Fisher, M. A., Plikaytis, B. B., & Shinnick, T. M. (2002). Microarray analysis of the *Mycobacterium tuberculosis* transcriptional response to the acidic conditions found in phagosomes. Journal of Bacteriology, 184(14), 4025–4032.

19. Garrett, C. K., Broadwell, L. J., Hayne, C. K., & Neher, S. B. (2015). Modulation of the activity of *Mycobacterium tuberculosis* LipY by Its PE Domain. PLoS One, 10(8), e0135447.

20. Ghazaei, C. (2018). *Mycobacterium tuberculosis* and lipids: Insights into molecular mechanisms from persistence to virulence. Journal of Research in Medical Sciences, 23, 63.

21. Griffin, J. E., Gawronski, J. D., Dejesus, M. A., Ioerger, T. R., Akerley, B. J., & Sassetti, C. M. (2011). High-resolution phenotypic profiling defines genes essential for mycobacterial growth and cholesterol catabolism. PLoS Pathogens, 7(9), e1002251.

22. Harding, C., & Boom, W. H. (2010). Regulation of antigen presentation by *Mycobacterium tuberculosis*: A role for Toll-like receptors. Nature Reviews Microbiology, 8(4), 296–307.

23. Hooft, R. W., Sander, C., & Vriend, G. (1997). Objectively judging the quality of a protein structure from a Ramachandran plot. Bioinformatics, 13(4), 425–430.

24. Humphrey, W., Dalke, A., & Schulten, K. (1996). VMD: visual molecular dynamics. Journal of Molecular Graphics, 14(1), 33–38.

25. Jones, D. T., & Swindells, M. B. (2002). Getting the most from PSI–BLAST. Trends in Biochemical Sciences, 27(3), 161–164.

26. Kumar A, Sharma A, Kaur G, Makkar P, Kaur J. Functional characterization of hypothetical proteins of Mycobacterium tuberculosis with possible esterase/lipase signature: a cumulative in silico and in vitro approach. J Biomol Struct Dyn. 2017 May;35(6):1226–1243.

27. Li, C., Li, Q., Zhang, Y., Gong, Z., Ren, S., Li, P., & Xie, J. (2017). Characterization and function of *Mycobacterium tuberculosis* H37Rv lipase Rv1076 (LipU). Microbiological Research, 196, 7–16.

28. Lin H, Xing J, Wang H, Wang S, Fang R, Li X, Li Z, Song N. Roles of Lipolytic enzymes in Mycobacterium tuberculosis pathogenesis. Front Microbiol. 2024 Jan 29;15:1329715.

29. MacGilvary, N. J., Kevorkian, Y. L., & Tan, S. (2019). Potassium response and homeostasis in *Mycobacterium tuberculosis* modulates environmental adaptation and is important for host colonization. PLoS Pathogens, 15(2), e1007591.

30. Mirdita M, Schütze K, Moriwaki Y, Heo L, Ovchinnikov S, Steinegger M. ColabFold: making protein folding accessible to all. Nat Methods. 2022 Jun;19(6):679–682.

31. Murry, J. P., Pandey, A. K., Sassetti, C. M., & Rubin, E. J. (2009). Phthiocerol dimycocerosate transport is required for resisting interferon-gamma-independent immunity. Journal of Infectious Diseases, 200, 774–782.

32. Murzin, A. G., Brenner, S. E., Hubbard, T., & Chothia, C. (1995). SCOP: a structural classification of proteins database for the investigation of sequences and structures. Journal of Molecular Biology, 247(4), 536–540.

33. Ollis DL, Cheah E, Cygler M, Dijkstra B, Frolow F, Franken SM, Harel M, Remington SJ, Silman I, Schrag J, et al. The alpha/beta hydrolase fold. Protein Eng. 1992 Apr;5(3):197–211.

34. Pettersen EF, Goddard TD, Huang CC, Couch GS, Greenblatt DM, Meng EC, Ferrin TE. UCSF Chimera--a visualization system for exploratory research and analysis. J Comput Chem. 2004 Oct;25(13):1605–12.

35. Pieper, U., Eswar, N., Stuart, A. C., Ilyin, V. A., & Sali, A. (2002). MODBASE, a database of annotated comparative protein structure models. Nucleic Acids Research, 30(1), 255–259.

36. Quigley, J., Hughitt, V. K., Velikovsky, C. A., Mariuzza, R. A., El-Sayed, N. M., & Briken, V. (2017). The cell wall lipid PDIM contributes to phagosomal escape and host cell exit of *Mycobacterium tuberculosis*. mBio, 8(3), e00148–17.

37. Ramachandran, G. N., & Sasisekharan, V. (1968). Conformation of polypeptides and proteins. Advances in Protein Chemistry, 23, 283–438.

38. Rameshwaram NR, Singh P, Ghosh S, Mukhopadhyay S. Lipid metabolism and intracellular bacterial virulence: key to next-generation therapeutics. Future Microbiol. 2018 Sep;13:1301–1328.

39. Rastogi, S., Singh, A. K., Pant, G., Mitra, K., Sashidhara, K. V., & Krishnan, M. Y. (2017). Down-regulation of PE11, a cell wall associated esterase, enhances the biofilm growth of *Mycobacterium tuberculosis* and reduces cell wall virulence lipid levels. Microbiology, 163(1), 52–61.

40. Reed, M. B., Domenech, P., Manca, C., Su, H., Barczak, A. K., Kreiswirth, B. N., … & Barry, C. E. 3rd. (2004). A glycolipid of hypervirulent tuberculosis strains that inhibits the innate immune response. Nature, 431(7004), 84–87.

41. Rustad, T. R., Harrell, M. I., Liao, R., & Sherman, D. R. (2008). The enduring hypoxic response of *Mycobacterium tuberculosis*. PloS One, 3(1), e1502.

42. Sampson, S. L. (2011). Mycobacterial PE/PPE proteins at the host-pathogen interface. Clinical and Developmental Immunology, 497203.

43. Sánchez, R., & Šali, A. (1997). Comparative protein structure modeling as an optimization problem. Journal of Molecular Structure: THEOCHEM, 398, 489–496.

44. Santucci, P., Smichi, N., Diomandé, S., Poncin, I., Point, V., Gaussier, H., … & Canaan, S. (2019). Dissecting the membrane lipid binding properties and lipase activity of *Mycobacterium tuberculosis* LipY domains. FEBS Journal, 286(16), 3164–3181.

45. Shen MY, Sali A. Statistical potential for assessment and prediction of protein structures. Protein Sci. 2006 Nov;15(11):2507–24.

46. Srivastava S, Battu MB, Khan MZ, Nandicoori VK, Mukhopadhyay S. Mycobacterium tuberculosis PPE2 Protein Interacts with p67phox and Inhibits Reactive Oxygen Species Production. J Immunol. 2019 Sep 1;203(5):1218–1229.

47. Silva, CAME., Rojony, R., Bermudez, LE., & Danelishvili, L. (2020). Short-Chain Fatty Acids Promote Mycobacterium avium subsp. hominissuis Growth in Nutrient-Limited Environments and Influence Susceptibility to Antibiotics. Pathogens, 9(9), 700.

48. Singh, G., Arya, S., Kumar, S., Narang, D., & Kaur, J. (2014). Molecular characterization of oxidative stress-inducible LipD of *Mycobacterium tuberculosis* H37Rv. Current Microbiology, 68(3), 387–396.

49. Singh, P., Rao, RN., Reddy, JR., Prasad, RB., Kotturu, SK., Ghosh, S., & Mukhopadhyay, S. (2016). PE11, a PE/PPE family protein of *Mycobacterium tuberculosis* is involved in cell wall remodeling and virulence. Scientific Reports, 6(1), 21624.

50. Sultana, R., Tanneeru, K., & Guruprasad, L. (2011). The PE-PPE domain in *Mycobacterium* reveals a serine α/β hydrolase fold and function: an in-silico analysis. PloS One, 6(2), e16745.

51. Sultana, R., Vemula, MH., Banerjee, S., & Guruprasad, L. (2013). The PE16 (Rv1430) of *Mycobacterium tuberculosis* is an esterase belonging to the serine hydrolase superfamily of proteins. PLoS One, 8(2), e55320.

52. Vandal OH, Nathan CF, Ehrt S. Acid resistance in Mycobacterium tuberculosis. J Bacteriol. 2009 Aug;191(15):4714–21.

53. van Kessel JC, Hatfull GF. Recombineering in Mycobacterium tuberculosis. Nat Methods. 2007 Feb;4(2):147–52.

54. Webb, B., & Sali, A. (2017). Protein structure modeling with MODELLER. In Functional Genomics: Methods and Protocols, 39-54.

55. Webb, B., & Sali, A. (2021). Protein Structure Modeling with MODELLER. Methods in Molecular Biology, 2199, 239–255.

56. Yang D, He X, Li S, Liu J, Stabenow J, Zalduondo L, White S, Kong Y. Rv1075c of Mycobacterium tuberculosis is a GDSL-Like Esterase and Is Important for Intracellular Survival. J Infect Dis. 2019 Jul 19;220(4):677-686.

57. Yang, D., Li, S., Stabenow, J., Zalduondo, L., & Kong, Y. (2019). *Mycobacterium tuberculosis* LipE has a Lipase/Esterase Activity and Is Important for Intracellular Growth and *In Vivo* Infection. Infection and Immunity, 88(1), e00750-19.

